# An aberrant cytoplasmic intron retention programme is a blueprint for ALS-related RBP mislocalization

**DOI:** 10.1101/2020.07.20.211557

**Authors:** Giulia E. Tyzack, Jacob Neeves, Pierre Klein, Hamish Crerar, Oliver Ziff, Doaa M. Taha, Raphaëlle Luisier, Nicholas M Luscombe, Rickie Patani

## Abstract

We recently described aberrant cytoplasmic SFPQ intron-retaining transcripts (IRTs) and concurrent SFPQ protein mislocalization as a new hallmark of amyotrophic lateral sclerosis (ALS). However the generalizability and potential roles of cytoplasmic IRTs in health and disease remain unclear. Here, using time-resolved deep-sequencing of nuclear and cytoplasmic fractions of hiPSCs undergoing motor neurogenesis, we reveal that ALS-causing VCP gene mutations lead to compartment-specific aberrant accumulation of IRTs. Specifically, we identify >100 IRTs with increased cytoplasmic (but not nuclear) abundance in ALS samples. Furthermore, these aberrant cytoplasmic IRTs possess sequence-specific attributes and differential predicted binding affinity to RNA binding proteins (RBPs). Remarkably, TDP-43, SFPQ and FUS – RBPs known for nuclear-to-cytoplasmic mislocalization in ALS – avidly and specifically bind to this aberrant cytoplasmic pool of IRTs, as opposed to any individual IRT. Our data are therefore consistent with a novel role for cytoplasmic IRTs in regulating compartment-specific protein abundance. This study provides new molecular insight into potential pathomechanisms underlying ALS and highlights aberrant cytoplasmic IRTs as potential therapeutic targets.

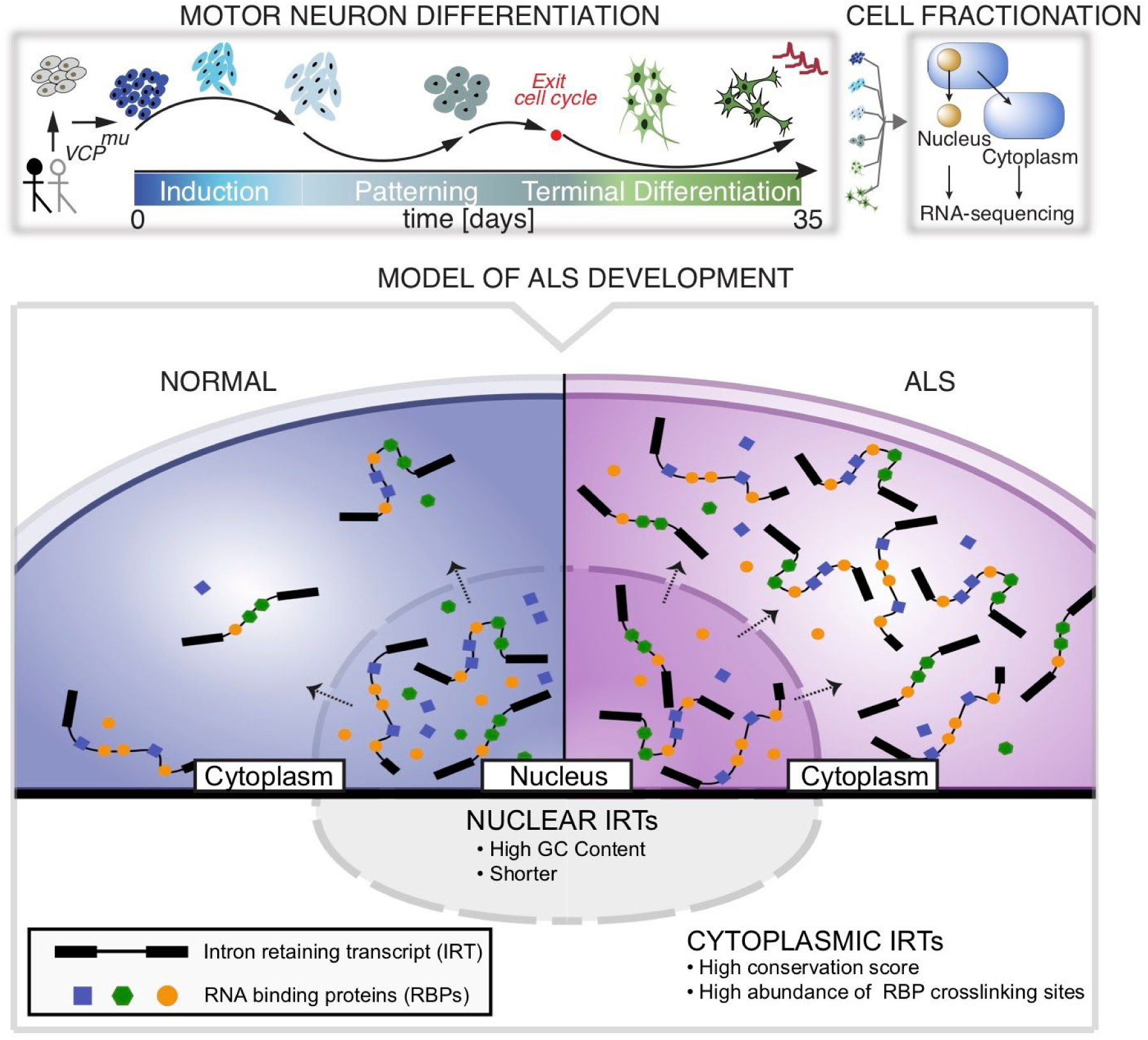

---

Studies have demonstrated that intron retention (IR) is more frequent in mammals than originally recognized, affecting transcripts from a majority of genes (Yap *et al*., 2012*a*, Braunschweig *et al*., 2014*a*; Boutz *et al*., 2015, Mauger *et al*., 2016*a*). Indeed, IR is a prominent mode of splicing during early neuronal differentiation (Braunschweig *et al*., 2014*a*; Luisier *et al*., 2018) and plays a functional role in neuronal homeostasis (Buckley *et al*., 2011*a*, Yap *et al*., 2012*b*, Braunschweig *et al*., 2014*a*, Mauger *et al*., 2016*b*). IR during neurogenesis has been shown to exhibit a set of features, an ‘‘IR code’’, that can reliably discriminate retained from constitutively spliced introns (Braunschweig *et al*., 2014*a*). IR has previously been implicated in regulating the transcriptome by coupling to RNA degradation pathways (Yap *et al*., 2012*a*; Colak *et al*., 2013; Wong *et al*., 2013, Braunschweig *et al*., 2014*a*; Kilchert *et al*., 2015). Although intron-retaining transcripts (IRTs) have predominantly been identified as residing within the nucleus where they are degraded (Yap *et al*., 2012*b*, Braunschweig *et al*., 2014*a*; Boutz *et al*., 2015), there is an expanding body of evidence demonstrating the cytoplasmic localisation of stable IRTs (Buckley *et al*., 2011*b*; Khaladkar *et al*., 2013; Sharangdhar *et al*., 2017; Saini *et al*., 2019; Price *et al*., 2020). It is noteworthy that neural cells, with their exceptional polarity and compartmentalisation, exhibit higher degrees of IR compared with other tissues (Yap *et al*., 2012*a*, Braunschweig *et al*., 2014*a*, Mauger *et al*., 2016*a*). However, their prevalence and role remain understudied. One of the few studies focusing on a cytoplasmic IRT showed that a retained intron in the Calm3 transcript determined its dendritic localization (Sharangdhar *et al*., 2017), thus revealing an addressing (or ‘zip-coding’) function of cytoplasmic IR. This study raises the possibility of new roles for intronic RNA sequences beyond a nuclear function, and suggests that cytoplasmic IR programmes are relevant to human neurological function and their perturbation, therefore, to disease.

Amyotrophic lateral sclerosis (ALS) is a rapidly progressive and incurable adult-onset condition, which leads to the relatively selective degeneration of motor neurons (MNs). The molecular pathological hallmark of ALS is the nuclear-to-cytoplasmic mislocalization of key RBPs (Neumann *et al*., 2006; Luisier *et al*., 2018; Tyzack *et al*., 2019), although the underlying mechanism remains elusive. ALS-causing gene mutations implicate crucial regulators of RNA-processing, which are normally expressed throughout development (Sreedharan *et al*., 2008; Vance *et al*., 2009). This raises the hypothesis that post-transcriptional changes, including those occurring during neurodevelopment, may play a pivotal role in the underlying molecular pathogenesis of ALS. We recently described IR as the predominant splicing event characterizing early stages of motor neuron lineage restriction from human iPSCs, which is perturbed by genetically diverse ALS-causing mutations (Luisier *et al*., 2018). However, whether this process affects the nuclear or cytoplasmic subcellular compartments similarly remains unresolved. Few studies have examined compartment-specific IR in differentiated neurons (Buckley *et al*., 2011*b*; Price *et al*., 2020), and to our knowledge, no study has comprehensively investigated cytoplasmic IR programmes during human motor neurogenesis, nor systematically characterized the effect of an ALS-causing mutation.

Here, we combine cellular fractionation of hiPSCs undergoing motor neurogenesis with deep RNA-sequencing (RNA-seq) of approximately 100 million paired end reads per sample to gain insights into the molecular ‘logic’ governing IR programmes in healthy and disease states. This is a rich resource for researchers across the disciplines of basic and applied neuroscience, constituting six timepoints during motor neuron neurogenesis for three control and three ALS hiPSCs lines, which have been separated into nuclear and cytoplasmic fractions.. Indeed, this resource allowed us to make important insights into the nature of aberrant IR in a human stem cell model of ALS. Specifically, we provide a taxonomy for aberrant IR based on nucleocytoplasmic distribution, *cis* attributes and predicted intron binding affinities to major RBPs. Consequently, we have uncovered a novel class of cytoplasmic IRTs that exhibits predictive value for the nuclear-to-cytoplasmic mislocalization of key RBPs, which constitutes a recognized molecular hallmark of ALS.

## RESULTS

### High coverage RNA-seq data from nuclear and cytoplasmic fractions during human motor neurogenesis

We analysed high-throughput poly(A) RNA-seq data derived from nuclear and cytoplasmic fractions of human induced pluripotent stem cells (hiPSC; day 0), neural precursors (NPC; day 3 and day 7), ‘patterned’ precursor motor neurons (ventral spinal cord; pMN; day 14), post-mitotic but electrophysiologically immature motor neurons (MN; day 22), and electrophysiologically mature MNs (mMNs; day 35). The cellular material was derived from two patients with the ALS-causing VCP gene mutation and four healthy controls (**Fig. 1A;** 95 samples from 6 time-points and 3 genotypes; 4 clones from 4 different healthy controls and 3 clones from 2 ALS patients with VCP mutations: R155C and R191Q, hereafter termed VCP^*mu*^). Cells from each stage of differentiation were characterised as previously reported (Hall *et al*., 2017). The efficiency of cellular fractionation was assessed both at protein and RNA level. The predominantly nuclear proteins histone H3 and PSPC1 were highly enriched in the nuclear fraction, whereas the cytosolic enzyme GAPDH was mainly detected in the cytoplasm (**Fig. 1B)**. Similarly, the presence of *GAPDH* intronic RNA was negligible in the cytoplasm, suggesting that leakage of RNA from the nucleus to the cytoplasm due to the fractionation protocol was minimal. Importantly, the efficiency of fractionation was comparable between control and VCP^*mu*^ lines (**Fig. 1B**). Singular value decomposition analysis of 18,834 reliably expressed genes across the 95 samples revealed that developmental stage and cellular fractions were the largest contributors to transcriptome diversity, explaining 41% and 15% of the variance, respectively (**Figs. 1C,D**). Notably the VCP^*mu*^ does not affect these processes and samples cluster with their age- and fraction-matched control counterparts (**Supplementary Fig. S1A**). Unsupervised hierarchical clustering (Spearman rank correlation and complete-linkage clustering) of the 95 samples using 18,834 genes segregated samples by developmental stage rather than genetic background (**Supplementary Fig. S1B**).

**Figure 1.**
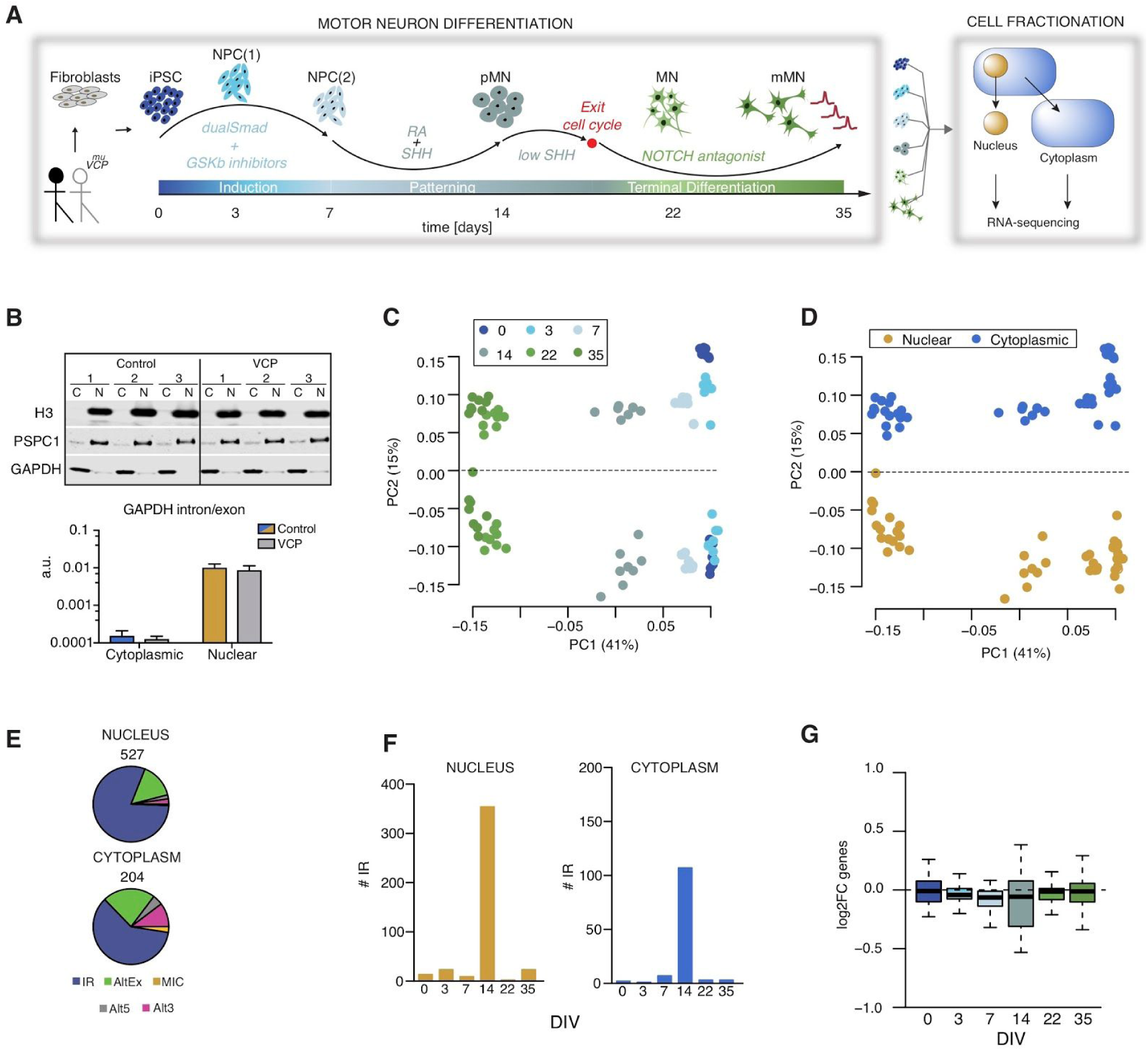
Time-resolved cellular fractionation and RNA-sequencing during human motor neurogenesis reveals widespread aberrant cytoplasmic IR in ALS. **A**. Schematic depicting the iPSC differentiation strategy for motor neurogenesis. Arrows indicate sampling time-points in days when cells were fractionated into nuclear and cytoplasmic compartments prior to deep (polyA) RNA-sequencing. Four iPSC clones were obtained from four different healthy controls and three iPSC clones from two ALS patients with VCP mutations: R155C and R191Q; hereafter termed VCP^*mu*^. Induced-pluripotent stem cells (iPSC); neural precursors (NPC); “patterned” precursor motor neurons (ventral spinal cord; pMN); post-mitotic but electrophysiologically inactive motor neurons (MN); electrophysiologically active MNs (mMN). **B**. Representative QC data for fractionation of samples at DIV=14 at protein level (Western blot, top) and RNA level (qPCR, bottom). In the Western blot, histone H3 and PSPC1 were chosen as protein markers for the nuclear faction, and GAPDH was used as cytosolic marker. In the qPCR, the ratio between intronic and exonic GAPDH sequences was measured in both fractions to exclude the leakage of nuclear RNA into the cytosolic fraction due to disruption of nuclei during the fractionation. Data is expressed as mean±SD from four lines per group. **C**. SVD performed on normalized 18,834 gene expression values across 95 samples. Samples are plotted by their coordinates along PC1 (41% of variance) and PC2 (15% of variance). Colors of data points indicate similar time in culture: iPSC (dark blue), DIV=3 (blue; NPC1), DIV=7 (light blue; NPC2), DIV=14 (grey; pMN), DIV=22 (lightgreen; MN) and DIV=35 (dark green; eMN). **D**. Same as (**C)** with colors of data points indicating similar cellular fractions: nuclear fraction (gold) and cytoplasmic fraction (blue). **E**. Pie charts representing proportions of included splicing events in VCP^*mu*^ at all timepoints of motor neurogenesis compared with age-matched control samples in nuclear (upper chart) and cytoplasmic (lower chart) fractions. Total number of events are indicated above the chart. Intron retention (IR); alternative exon (AltEx); microexons (MIC); alternative 5′ and 3′ UTR (Alt5 and Alt3). **F**. Bar graphs representing the number of retained introns in VCP^*mu*^ compared to control samples at specific timepoints during MN differentiation. Nuclear fraction (gold; left). Cytoplasmic fraction (blue; right). **G**. Boxplots showing the distributions of cytoplasmic log2 fold-changes for 72 essential splicing factor genes (**Table S8**) between VCP^*mu*^ and controls.

### Widespread aberrant cytoplasmic IR in a human stem cell model of ALS

We previously identified ALS-related aberrant cytoplasmic SFPQ IRT and concurrent protein mislocalization (Luisier *et al*., 2018). Here we tested whether aberrant cytoplasmic IR is a generalizable transcriptomic phenomenon in ALS. We examined patterns of splicing using the RNA-seq pipeline VAST-TOOLS (Irimia *et al*., 2014). We identified 791 nuclear (527 included and 264 skipped) and 329 cytoplasmic (204 included and 125 skipped) alternative splicing events that are statistically significantly different between VCP^*mu*^ and control samples in at least one time-point (**Supplementary Fig. 1G,H**). In line with our previous study, the majority of inclusion events between VCP^*mu*^ and control samples were retained introns (**Fig. 1E**). The VCP mutation not only leads to increased IR in the nuclear compartment but also in the cytoplasm (**Supplementary Figs. S1C-F**). We find that these events peak in pMNs (day 14 *in vitro*) (**Fig. 1F**) when we observe a coincident decrease in expression of splicing factors (**Fig. 1G** and **Supplementary Fig. S1I**); most notable is the 112 aberrant IR events in the cytoplasmic fraction. Collectively, these findings demonstrate that aberrant cytoplasmic IR is a widespread phenomenon in ALS that occurs during motor neurogenesis.

### A nucleocytoplasmic taxonomy for aberrant IRTs

We next manually curated the list of nuclear and cytoplasmic VCP-mutation related aberrant IR events, focusing on pMNs (day 14 in vitro). We identified three categories of introns that are aberrantly retained in VCP^*mu*^ : 1) 237 predominantly in the nucleus; 2) 63 in both the nucleus and the cytoplasm; 3) 49 predominantly in the cytoplasm (**Fig. 2A, Tables S3-S5**). Gene Ontology functional enrichment analysis shows the specific biological association of affected genes, including cell cycle for the predominantly nuclear IRTs and protein localisation for those that are predominantly cytoplasmic (**Fig. 2B**). We previously identified aberrant IR in *SFPQ, FUS* and *DDX39A* transcripts in VCP^*mu*^ at an early stage of MN development (Luisier *et al*., 2018), which we validated here by RT-PCR. Importantly these three IR events belong to the third category (**Fig. 2C-D; Supplementary Fig. 2A-B**). The finding that a large number of aberrantly retained introns, including SFPQ, DDX39A and FUS, are predominantly localized within the cytoplasm suggests nuclear export and/or cytoplasmic stabilisation, further supporting the hypothesis that VCP^*mu*^ impacts distinct IRTs and may lead to stereotyped cellular dysfunction as a consequence.

**Figure 2.**
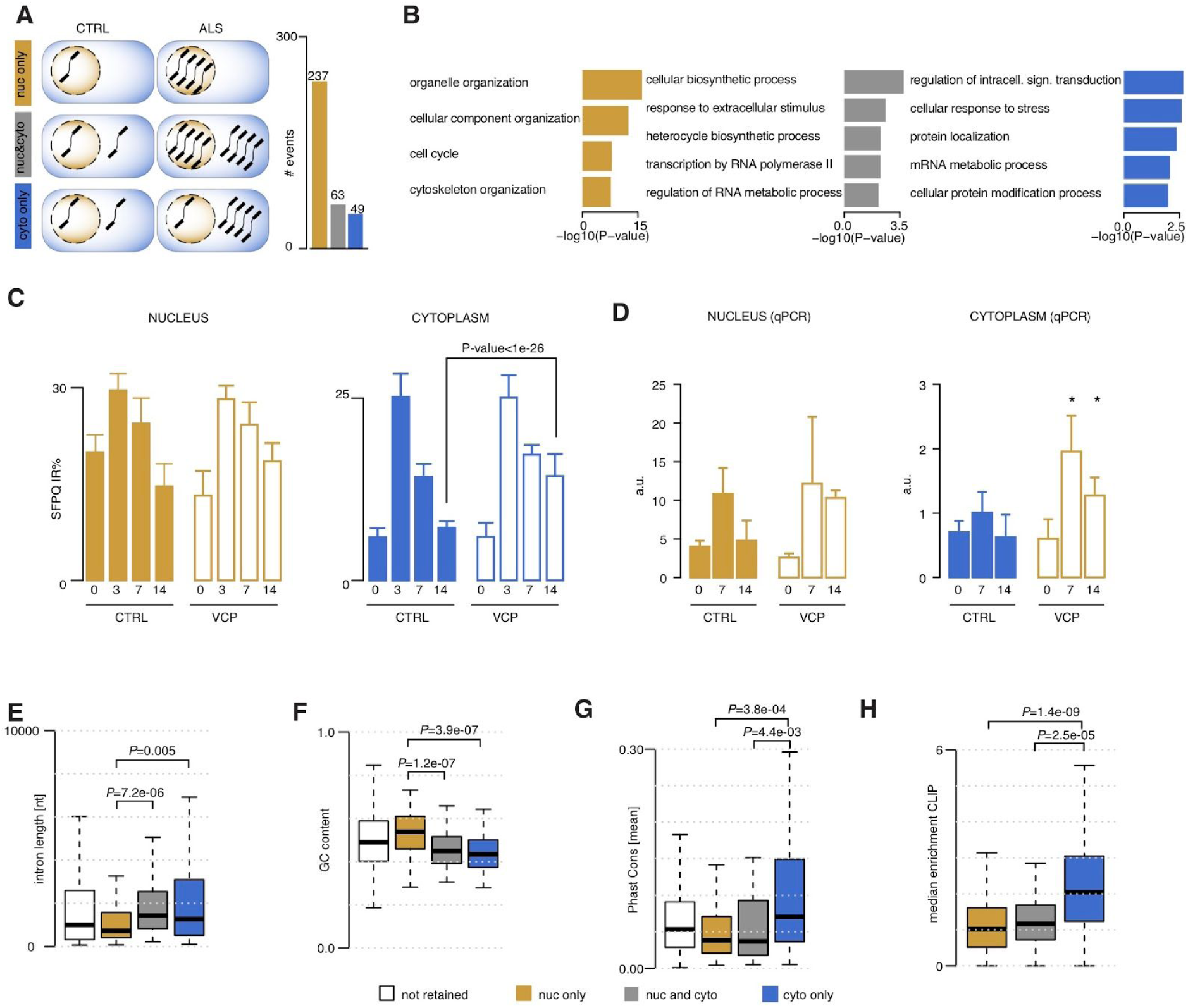
Aberrant nuclear and cytoplasmic intronic sequences exhibit distinct characteristics. **A**. Schematic of our proposed taxonomy for aberrant IRTs (left) and bar graphs (right) representing the numbers of retained introns in VCP^*mu*^ compared to control samples at DIV=14 that are predominantly nuclear (gold bar), aberrant in both nucleus and cytoplasm (grey bar), or predominant in cytoplasm (blue bar). The number of events in each category is indicated above the bar. **B**. Bar plots displaying the enrichment scores for GO biological functions of genes that are targeted by each group of aberrantly retained introns. **C**. Bar graphs quantifying percentage IR in SFPQ gene at DIV=0, 3, 7 and 14 in control and *VCP*_*mu*_ samples (mean ± s.d.; Fisher count test) in the nucleus (left) and cytoplasm (right). **D**. Bar plots displaying SFPQ IR levels measured in the nucleus (left) and cytoplasm (right) by qPCR at DIV=0, 7, 14 in control and *VCP*^*mu*^ samples (mean±s.d. from four lines per group, *p<0.05, individual t-tests comparing control and *VCP*^*mu*^ samples at each time point). **E, F, G, H**. Comparison of intron length, GC content (%), conservation scores and median enrichment for RBP binding sites of the three groups of aberrantly retained introns. Data shown as box plots in which the centre line is the median, limits are the interquartile range and whiskers are the minimum and maximum. P-values obtained from Mann-Withney test. All introns in the gene-set targeted by IR in VCP^*mu*^ at DIV=14 (white).

### Molecular characteristics of aberrantly retained introns

Prior studies have shown that retained introns are on average shorter and more G/C rich(Galante *et al*., 2004; Sakabe and de Souza, 2007, Braunschweig *et al*., 2014*a*). Strikingly, here we find that only the predominantly nuclear aberrantly retained introns exhibit those features. In complete contrast, aberrantly retained introns within the cytoplasm (in other words, those in both nucleus and cytoplasm, and cytoplasm only) are on average longer and have lower GC content (**Figs. 2E,F**). Furthermore, the predominantly nuclear aberrant IRTs correlate with a cytoplasmic decrease in gene expression of their non-intron-retaining counterparts: this is consistent with prior observations showing that nuclearly-detained IRTs reduce the level of gene expression (Braunschweig *et al*., 2014*a*). Conversely, IR transcripts found in the cytoplasm correlate with increased gene expression within the nucleus (**Supplementary Fig. 2E)**. Importantly this suggests that previously reported features (Galante *et al*., 2004; Sakabe and de Souza, 2007, Braunschweig *et al*., 2014*a*) discriminate nuclearly detained IRTs from cytoplasmic ones. Two additional features further discriminate cytoplasmic-only events from those in both compartments: 1) a high conservation score (**Fig. 2G**) and 2) a greater abundance of RNA-binding protein (RBP) crosslinking events in the cytoplasmic only retained introns (Sloan *et al*., 2016; Van Nostrand *et al*., 2017) (**Fig. 2H**).

### Aberrant cytoplasmic IRTs blueprint ALS-related protein-mislocalization

The finding that cytoplasmic aberrant IRTs avidly bind to RBPs raises the hypothesis that this interaction drives hallmark RBP mislocalization events in ALS. Indeed we previously reported that the aberrant SFPQ IRT and the SFPQ protein itself, which are predicted to avidly bind to each other, are both exported to the cytoplasm thus providing a potential mechanism for SFPQ mislocalization in ALS (Luisier *et al*., 2018). We therefore further examined the nature of interaction between RBPs and aberrant cytoplasmic IRTs using our richer dataset. At least 27 RBPs systematically exhibit statistically significant increased binding to cytoplasmic-only retained introns compared with their predominantly nuclear counterparts (**Fig. 3A**). These RBPs form a densely connected network of experimentally validated interacting proteins that are enriched in mRNA metabolism functions (**Fig. 3B**). The network includes a subset of 9 RBPs with known functions in processing capped intron-containing pre-mRNA, which further implicates disrupted post-transcriptional splicing in ALS pathogenesis. Importantly, also within this network of RBPs are those that exhibit hallmark nuclear-to-cytoplasmic mislocalization in ALS: TDP-43 (**Fig. 3B**), SFPQ (**Fig. 3C**) and FUS (**Supplementary Fig. 2F)** (Neumann *et al*., 2006; Luisier *et al*., 2018; Tyzack *et al*., 2019). Taken together, these findings support the hypothesis that ALS leads to an increase in the cytoplasmic abundance of a class of RNA with a large capacity for binding proteins, thus creating an environment that might encourage mislocalization of their cognate RBPs **(Fig. 3D)**.

**Figure 3.**
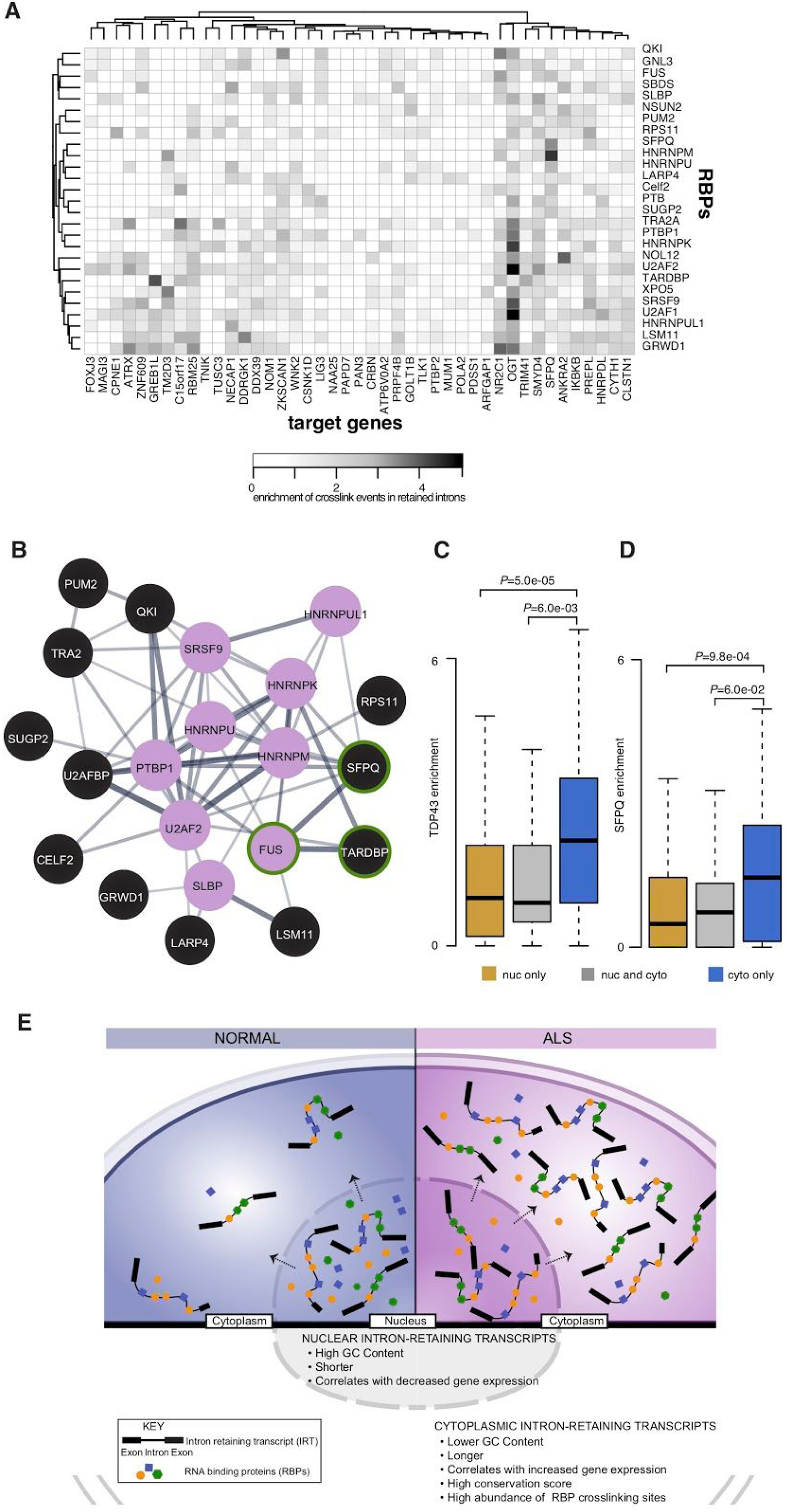
Cytoplasmic intron-retaining transcripts create a mislocalization-prone environment for bound RBPs. **A**. Heatmap of the enrichment score of the crosslinking events in each of the 49 predominantly cytoplasmic aberrant IRTs for 27 RBPs that exhibit significantly higher enrichment compared to the two other categories of IRTs (i.e. predominantly nuclear and those that are both cytoplasmic and nuclear). **B**. Network of protein–protein interactions for 21 (out of the 27) RBPs enriched in cytoplasmic aberrant retained introns. Edges represent experimentally determined protein–protein interactions annotated in the STRING database (Szklarczyk *et al*., 2017). 9 of these RBPs belong to the *Processing of Capped Intron-Containing Pre-mRNA* Reactome(Croft *et al*., 2014) pathway (filled magenta circles) and 3 are RBPs that exhibit hallmark nuclear-to-cytoplasmic mislocalization ALS (green circle). Line thickness indicates the strength of data support based on text mining and experiments. **C, D**. Comparison of enrichment across all genes within each category for TDP43 and SFPQ crosslinking events in the retained introns between the 3 groups of aberrantly retained introns. Data shown as box plots in which the centre line is the median, limits are the interquartile range and whiskers are the minimum and maximum. P-values obtained from Mann-Withney test. **E**. Schematic diagram of proposed model where cytoplasmic intron-retaining transcript accumulation in ALS leads to protein mislocalization.

## DISCUSSION

Recent studies have demonstrated that IR is more frequent in mammals than previously recognized (Yap *et al*., 2012*a*, Braunschweig *et al*., 2014*a*; Boutz *et al*., 2015, Mauger *et al*., 2016*a*). It is however noteworthy that the majority of studies have focused on nuclear IRTs and that the importance of cytoplasmic IRTs remains relatively understudied, particularly in the contexts of neuronal development and disease. We previously identified aberrant IR in the SFPQ transcript across human stem cell models of diverse genetic forms of ALS (including those caused by mutations in VCP, SOD1 and FUS genes) (Luisier *et al*., 2018). In the present study, we sought to understand the generalizability of cytoplasmically localised aberrant IRTs in ALS by combining directed differentiation of patient-specific hiPSCs into spinal MNs, with cellular fractionation and by deep (polyA) RNA sequencing. We showed that aberrant cytoplasmic IR is indeed a widespread molecular phenomenon in ALS that comprises at least 112 transcripts including SFPQ. To better understand the nature of cytoplasmic IRTs, we categorized the aberrant IR events into three classes according to their nucleocytoplasmic distribution (nuclear, cytoplasmic and those present in both compartments) and examined their cis attributes.

Retained introns during neurogenesis have previously been shown to exhibit a highly correlated set of *cis* features comprising an ‘‘IR code’’ that can reliably discriminate retained from constitutively spliced introns (Braunschweig *et al*., 2014*a*). Notably the aberrantly retained introns that are predominantly in the nucleus of VCP mutant cultures display similar lengths and GC content to this ‘IR code’(Braunschweig *et al*., 2014*a*). IR has been previously implicated in fine-tuning the cellular transcriptome by targeting transcripts to RNA degradation pathways such as nonsense mediated decay (Yap *et al*., 2012*a*; Colak *et al*., 2013; Wong *et al*., 2013, Braunschweig *et al*., 2014*a*; Kilchert *et al*., 2015). The retained introns characterised in the aforementioned studies act broadly to reduce the levels of transcripts that are not required (Boutz *et al*., 2015; Braun *et al*., 2017). Indeed only this category of aberrant IR correlates with reduced gene expression in our data. Remarkably, the two other categories we identified exhibit an almost opposite effect on their gene expression levels and thus stimulated further investigation. Our study further revealed a specific class of cytoplasmic IR transcripts that 1) have unique features compared to those reported in previous studies and 2) have conspicuously high affinity for RBPs, including those that are mislocalized in ALS (TDP43, FUS and SFPQ)(Neumann *et al*., 2006; Luisier *et al*., 2018; Tyzack *et al*., 2019). These findings raise the hypothesis that a subset of cytoplasmic IRTs have a distinct role compared to the previously reported IRTs that regulate gene expression and translation through coupling with nonsense mediated decay (NMD)(Braunschweig *et al*., 2014*a*). Indeed RNA localization to distinct subcellular compartments has been shown to regulate spatio-temporal control of protein expression(Holt and Bullock, 2009) but less is known about their role in protein localisation. Here we propose that a subset of IRTs aberrantly accumulate in the cytoplasm and their intronic sequences serve as ‘blueprints’ for the hallmark protein mislocalization events in ALS by creating a mislocalization-prone environment for their bound (and usually predominantly nuclear) RBPs (**Fig. 3E**). This is reinforced by the fact that the RBPs with the largest difference in binding affinity between the predominantly cytoplasmic vs predominantly nuclear aberrant IR are those known to be mislocalized in ALS: TDP-43, FUS and SFPQ. Future studies will directly assess this hypothesis. In summary, we propose that cytoplasmic retained introns function as RNA regulators in the homeostatic control of RBP localisation and that an ALS-related aberrant increase in cytoplasmic IR transcripts disrupts this process.

---

**Supplementary Figure 1.**
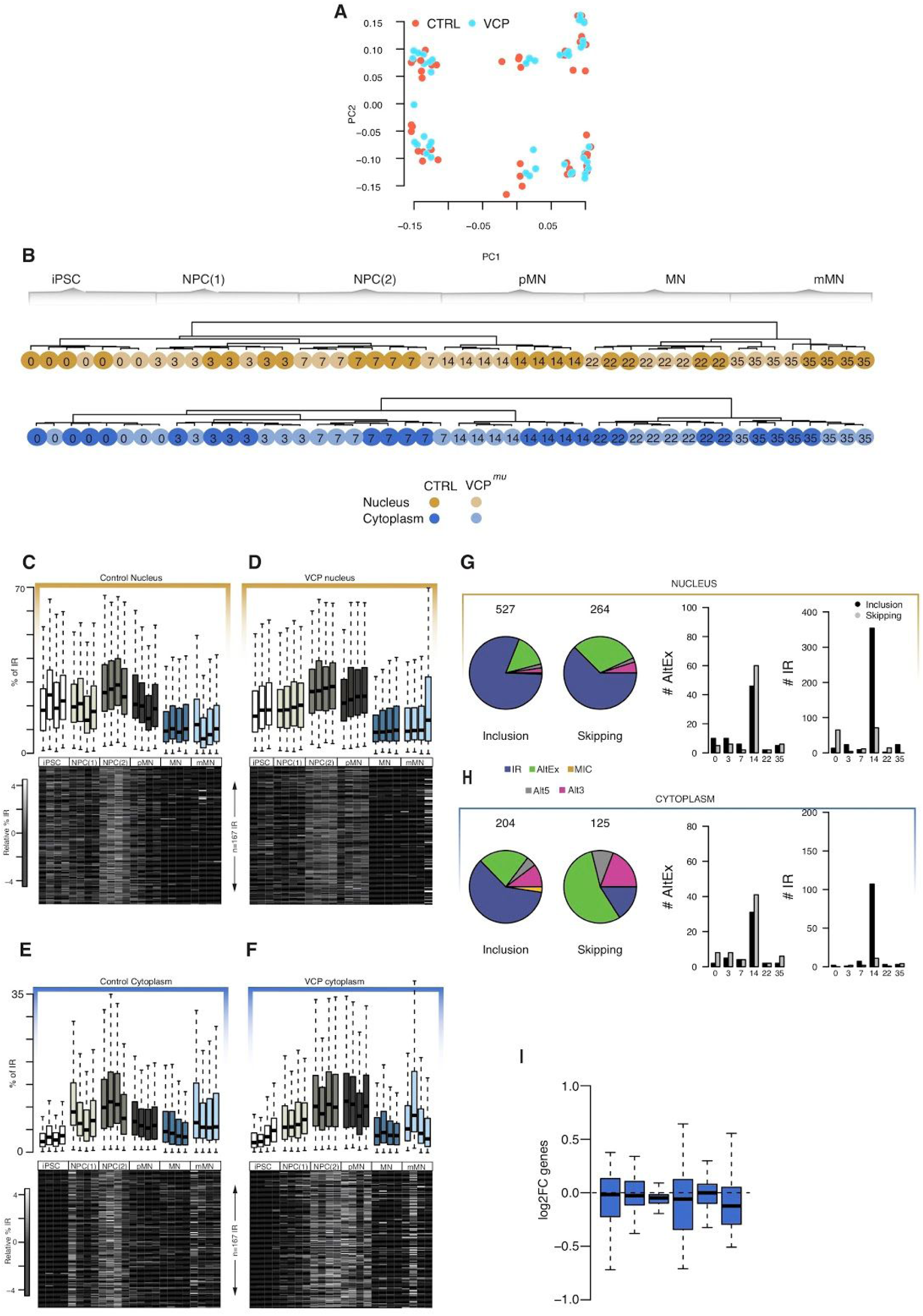
**A**. SVD performed on normalized 18,834 gene expression values across 95 samples. Samples are plotted by their coordinates along PC1 (41% of variance) and PC2 (15% of variance). Colors of data points indicate control or mutated samples: control (red), VCP^*mu*^ (magenta). **B**. Unsupervised hierarchical clustering of 18,834 genes groups the 47 nuclear samples (upper) and 48 cytoplasmic samples (lower) according to developmental stage, rather than genetic background. Dark gold circles = nuclear control samples; light gold circles = nuclear VCP^*mu*^ samples; Dark blue circles =cytoplasmic control samples; light blue circles =cytoplasmic VCP^*mu*^ samples; sampling time-points are indicated inside the circles. **C, D, E, F**. Heatmaps of the standardised relative percentage of IR in 167 introns identified in (Luisier *et al*., 2018) in replicate samples at each differentiation stage in nuclear control samples, nuclear VCP^*mu*^ samples, cytoplasmic control samples, cytoplasmic VCP^*mu*^ samples respectively. **G, H**. (left) Pie charts representing proportions of all included and skipped splicing events in VCP^*mu*^ at any stages of motor neurogenesis compared with age-matched control samples in nuclear and cytoplasmic fractions respectively. Total numbers of events are indicated above the chart. Intron retention (IR); alternative exon (AltEx); microexons (MIC); alternative 5′ and 3′ UTR (Alt5 and Alt3). (right) Bar graphs representing the numbers of exonic (black bars) and intronic (grey bars) splicing events, in VCP^*mu*^ compared to control samples at specific timepoints during MN differentiation. **I**. Boxplots showing the distributions of nuclear log2 fold-changes for 72 essential splicing factor genes (**Table S8**) between VCP^*mu*^ and controls. Data shown as box plots in which the centre line is the median, limits are the interquartile range and whiskers are the minimum and maximum.

**Supplementary Figure 2.**
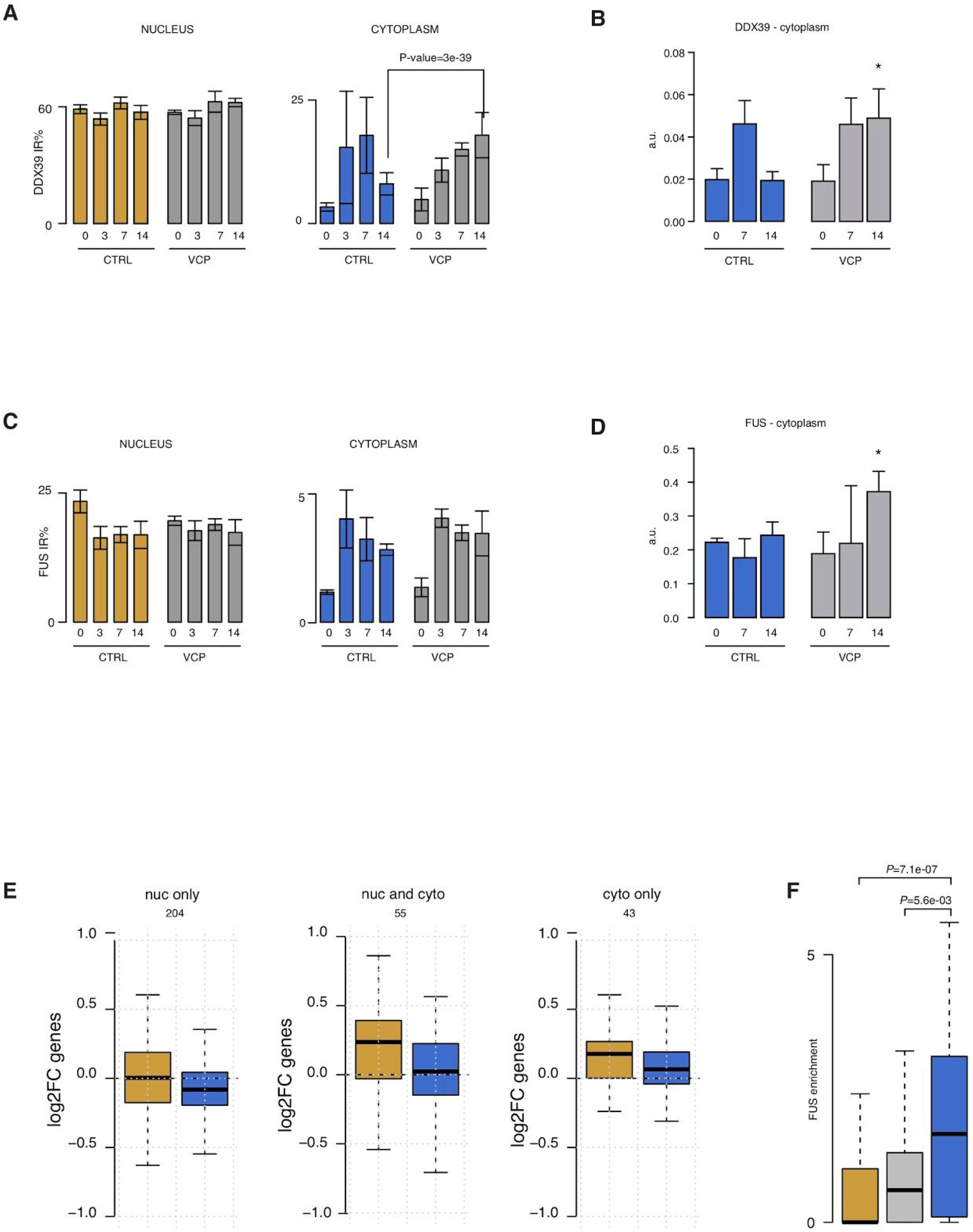
**A**. Bar graphs quantifying percentage IR in DDX39A gene at DIV=0, 3, 7 and 14 in control and *VCP*^*mu*^ samples (mean ± s.d.; Fisher count test) in the nucleus (left) and cytoplasm (right). **B**. Bar plots displaying DDX39A IR levels measured in the cytoplasm by qPCR at DIV=0, 7, 14 in control and *VCP*^*mu*^ samples (mean±s.d. from 4 lines per group, *p<0.05, individual t-tests comparing control and *VCP*^*mu*^ samples at each time point). **C**. Same as A but for FUS gene. **D**. Same as B but for FUS gene. **E**. Boxplots showing the distributions of nuclear (gold) and cytoplasmic (blue) log2 fold-changes for 204, 55 and 43 genes belonging to the three groups of aberrant IR between VCP^*mu*^ and controls. Data shown as box plots in which the centre line is the median, limits are the interquartile range and whiskers are the minimum and maximum. **F**. Comparison of enrichment for FUS crosslinking events between the 3 groups of aberrantly retained introns. Predominantly nuclear (gold), nuclear and cytoplasmic (grey), and predominantly cytoplasmic (blue). Data shown as box plots in which the centre line is the median, limits are the interquartile range and whiskers are the minimum and maximum. P-values obtained from Mann-Withney test.

## AUTHOR CONTRIBUTIONS

Conceptualization, R.L., N.M.L., R.P.; Formal Analysis, R.L.; Investigation, R.L., G.E.T., J.N., H.C., P.K., O.Z. D.M.T.; Writing – Original Draft, R.L., R.P.; Writing – Review & Editing, R.L., N.M.L., R.P, G.E.T, J.N.; Resources, R.L., N.M.L., R.P.; Visualization, R.L.; Funding Acquisition, R.L., N.M.L., R.P.; Supervision, R.L, N.M.L., R.P.

## ACKNOWLEDGMENTS

The authors wish to thank the patients for fibroblast donations. We also thank Anob M Chakrabarti for sharing BED files of aligned CLIP data. We thank Mahmoud-Reza Rafiee for useful discussions and advice regarding library preparation for RNA sequencing. We are grateful for the help and support provided by the Scientific Computing section and the DNA Sequencing section of Research Support Division at OIST. This work was supported by the Idiap Research Institute and by the Francis Crick Institute which receives its core funding from Cancer Research UK (FC010110), the UK Medical Research Council (FC010110), and the Wellcome Trust (FC010110). D.M.T. is supported by a Newton-Mosharafa scholarship. R.P. holds an MRC Senior Clinical Fellowship [MR/S006591/1]. We also acknowledge Wellcome Trust funding to N.M.L.[103760/Z/14/Z], an MRC eMedLab Medical Bioinformatics Infrastructure Award to N.M.L. (MR/L016311/1). N.M.L. is a Winton Group Leader in recognition of the Winton Charitable Foundation’s support towards the establishment of the Francis Crick Institute.

## ONLINE METHODS

### Contact for reagent and resource sharing

Further information and requests for resources and reagents should be directed to and will be fulfilled by the Lead Contact, Rickie Patani.

### Ethics Statement

Informed consent was obtained from all patients and healthy controls in this study. Experimental protocols were all carried out according to approved regulations and guidelines by UCLH’s National Hospital for Neurology and Neurosurgery and UCL’s Institute of Neurology joint research ethics committee (09/0272).

### Cell Culture

Induced PSCs were maintained on Geltrex (Life Technologies) with Essential 8 Medium media (Life Technologies), and passaged using EDTA (Life Technologies, 0.5mM). All cell cultures were maintained at 37C and 5% carbon dioxide. Details of the lines used in this study are provided in **Table S1**. One of the control lines used (control 3) is commercially available and was purchased fromThermoFisher Scientific (cat. number A18945).

### Motor neuron differentiation

Motor neuron (MN) differentiation was carried out using an adapted version of a previously published protocol (Hall *et al*., 2017). Briefly, iPSCs were first differentiated to neuroepithelium by plating on Geltrex-coated plates to 100% confluency in chemically defined medium consisting of DMEM/F12 Glutamax, Neurobasal, L-Glutamine, N2 supplement, non essential amino acids, B27 supplement, β-mercaptoethanol (all from Life Technologies) and insulin (Sigma). Treatment with small molecules from day 0-7 was as follows: 1µM Dorsomorphin (Millipore), 2µM SB431542 (Tocris Bioscience), and 3.3µM CHIR99021 (Miltenyi Biotec). Starting from day 8, the neuroepithelial layer was patterned for 7 days with 0.5µM retinoic acid and 1µM Purmorphamine. At day 14 pMN precursors were treated with 0.1µM Purmorphamine for a further 4 days before being terminally differentiated in 0.1 µM Compound E (Enzo Life Sciences) to promote cell cycle exit. At relevant time points (day 22 and day 35) cells were harvested for cellular fractionation.

### Cell fractionation

Biochemical subcellular fractionation was achieved for all cell stages using the Ambion PARIS kit (ThermoFisher Scientific) cell fractionation buffer, following the manufacturer’s general protocol, and an 8M Urea Nuclear Lysis Buffer prepared in house. Initially, cells were washed once using ice-cold PBS. Cytosolic fraction was then obtained by lysing cells directly in ice-cold cell fractionation buffer (ThermoFisher Scientific) for 3-15 minutes, thus disrupting plasma membranes, whilst leaving nuclear membranes intact. Lysates were then centrifuged for 3 minutes at 500 x g at 4C. The supernatant was then collected, further centrifuged at maximum speed in a bench centrifuge at 4C for 1 minute, and the resulting supernatant was then processed as cytosolic fraction. Nuclear pellets from the first centrifugation step were gently washed once with cell fractionation buffer and then lysed on ice for 30 minutes in 8M Urea Nuclear Lysis Buffer, containing 50mM Tris-HCL (pH 8), 100mM NaCl, 0.1% SDS, and 1mM DTT. The resulting nuclear fraction was then homogenised using a QIAshredder (QIAGEN) to shred chromatin and reduce viscosity, before being further processed for RNA extraction. Both lysis buffers were supplemented with 0.1 U/µl RiboLock RNase Inhibitor (ThermoFisher Scientific) and HALT Protease Inhibitor Complex (ThermoFisher Scientific).

### RNA extraction and sequencing

The Promega Maxwell RSC simplyRNA cells kit including DNase treatment, alongside the Maxwell RSC instrument, was used for RNA extractions. For qPCR validations, RNA was extracted using the RNeasy Plus Mini Kit (Qiagen). The nanodrop was used to assess RNA concentration and the 260/280 ratio, and the Agilent TapeStationwas used to assess quality. RNA integrity (RIN) scores were analysed to quality check all samples used in this work.

### RNA-sequencing data

Paired-end polyA stranded RNAseq libraries were prepared from fractionated nucleus and cytoplasm obtained from 6 distinct stages of motor neuron differentiation from control and VCP^*mu*^ samples (iPSC, and days 3, 7, 14, 21 and 35) using the NEBNext® Ultra™ II Directional RNA Library Prep Kit for Illumina®, with NEBNext® Poly(A) mRNA Magnetic Isolation Module with 500ng of total RNA as input. Libraries were sequenced at the OIST DNA sequencing section using the NovaSeq 6000 Sequencing technology. 50 bp-long reads were trimmed for adapter sequence and initially aligned to ribosomal RNA sequences to filter out reads that may come from ribosomal RNA contamination using bowtie2 (-v 0) (Langmead and Salzberg, 2012). The remaining reads were aligned to the human genome (hg38) using the splice aware aligner STAR (STAR-2.6.0)(Dobin *et al*., 2013) with default parameters. All libraries generated in this study had <1% rRNA, <1% mtDNA, >90% strandedness and >70% exonic reads (data not shown). One control iPSC nuclear sample failed quality control and was discarded. The list of samples together with the corresponding number of reads in each library and alignment statistics are provided in **Table S2**. All sequence data for this project has been deposited at NCBI GEO database under accession number **GSE152983**.

### Gene expression analysis

Kallisto (Bray *et al*., 2016) was used to (1) build a transcript index from the Gencode hg38 release Homo sapiens transcriptome (-k 31), (2) pseudo-align the RNA-seq reads to the transcriptome and (3) quantify transcript abundances (-b 100 -s 50—rf-stranded). Subsequent analysis was performed with the R statistical package version 3.3.1 (2016) and Bioconductor libraries version 3.3 (R Core Team. R: A Language and Environment for Statistical Computing. Vienna, Austria: R Foundation for Statistical Computing; 2013). Kallisto outputs transcript abundance, and thus we calculated the abundance of genes by summing up the estimated raw count of the constituent isoforms to obtain a single value per gene. For a given sample, the histogram of log2 gene count is generally bimodal, with the modes corresponding to non-expressed and expressed genes. Reliably expressed genes for each condition (VCP^*mu*^ or control at days 0, 3, 7, 14, 22 and 35 in each fraction) were next identified by fitting a two-component Gaussian mixture to the log2 estimated count gene data with R package mclust (Fraley and Raftery, n.d.); a pseudocount of 1 was added before log2 transformation. A gene was considered to be reliably expressed in a given condition if the probability of it belonging to the non-expressed class was under 1% in each sample belonging to the condition. 18,834 genes were selected based on their detected expression in at least one of the 24 conditions (i.e. 6 different timepoints of lineage restriction for control and VCP^*mu*^ in nuclear and cytoplasm). Next we quantile normalized the columns of the gene count matrix with R package limma (Boldstad *et al*., 2003). Unsupervised hierarchical clustering of the filtered and normalised gene count matrix was performed with Spearman rank correlation as a distance measure and complete clustering algorithm. Principal component analysis has been done with the svd function in R. Differential gene expression analysis was performed with Sleuth comparing VCP mutant or control at each day (0, 3, 7, 14, 22 and 35) in each fraction. Genes that showed a log twofold differential expression and a P-value < 0.05, and that were reliably expressed in either VCP mutant or control condition were considered as changing significantly.

### Splicing analysis

Intron retention (IR) focussed analysis has been performed on the 167 IR events previously found to be retained during MN differentiation(Luisier *et al*., 2018) for which a percentage of IR has been calculated as the fraction of intron mapping reads to the average number of reads mapping to the adjacent 5’ and 3’ exons normalised to the length of the respective intron and exons. A Fisher count test P-value has been obtained when testing for differential IR between conditions. Next the identification of all classes of alternative splicing (AS) events in motor neuron differentiation was performed with the RNA-seq pipeline *vast-tools* (Irimia *et al*., 2014). For an AS event to be considered differentially regulated between two conditions, we required a minimum average ΔPSI (between the paired replicates) of at least 15% and that the transcript targeted by the splicing event in question to be reliably expressed in all samples from the conditions compared i.e enough read coverage in all samples of interest. We next focussed on the introns aberrantly retained at dat 14 in VCP mutant compared to control samples and conducted Integrative Genomics Viewer (IGV)-guided manual curation to remove low coverage IR obtaining 237 high-confidence IR events that were aberrantly retained predominantly in the nucleus, 63 introns aberrantly retained in both the nucleus and the cytoplasm, and 49 introns predominantly in the cytoplasm. These and their associated characteristics (GC content, conservation score, enrichment in CLIP binding sites) are reported in **Tables S3-S6**.

### GO enrichment analysis

GO enrichment analysis was performed using classic Fisher test with topGO Bioconductor package(Alexa and Rahnenfuhrer, 2016). Only GO terms containing at least 10 annotated genes were considered. A P-value of 0.05 was used as the level of significance. On the figures, top significant GO terms were manually selected by removing redundant GO terms and terms which contain fewer than five significant genes.

### Mapping and analysis of CLIP data

To identify RBPs that bind to aberrantly retained introns, we examined iCLIP data for 21 RBPs(Attig *et al*., 2018), and eCLIP data from K562 and HepG2 cells for 112 RBPs available from ENCODE (Sloan *et al*., 2016; Van Nostrand *et al*., 2017). Before mapping the reads, adapter sequences were removed using Cutadapt v1.9.dev1 and reads shorter than 18 nucleotides were dropped from the analysis. Reads were mapped with STAR v2.4.0i (Dobin *et al*., 2013) to UCSC hg19/GRCh37 genome assembly. The results were lifted to hg38 using liftOver(Hinrichs *et al*., 2006) To quantify binding to individual loci, only uniquely mapping reads were used. Relative enrichment for each of the RBPs was obtained by computing the proportion of crosslink events mapping to retained intron compared to non-retained introns of the same genes.

### Reverse transcription, qPCR and intron retention validation

Reverse transcription was performed using the Revert Aid First Strand cDNA Synthesis Kit (ThermoFisher Scientific) using 500ng-1μg of total RNA and random hexamers. qPCR was performed using the PowerUP SYBR Green Master Mix (ThermoFisher Scientific) and the Agilent Mx3000P QPCR System or the QuantStudio 6 Flex Real-Time PCR System (Applied Biosystems). Primers used are listed in **Table S7**. Specific amplification was determined by melt curve analysis and agarose gel electrophoresis of the PCR products. Primer pairs with 90-110% efficiency were used. Intron retention validation was performed as previously described (Luisier *et al*., 2018), and RT-minus samples were used as negative controls. Data was analysed using the *ddCt* method and is expressed as arbitrary units (a.u. = 2^*-dCt*^).

### Electronic supplementary material

Supplementary Tables 1-8 can be accessed <u>here</u>.

Table S1 | Description of human sample origin and mutations.

Table S2 | Detailed results of the quality control of the 288 fastq files.

Table S3 | List of the 237 IR events that are aberrant at DIV=14 in VCP *nuclear* compartment predominantly.

Table S4 | List of the 63 IR events that are aberrant at DIV=14 in VCP *nuclear* and *cytoplasmic* compartments.

Table S5 | List of the 49 IR events that are aberrant at DIV=14 in VCP *cytoplasmic* compartment predominantly.

Table S6 | Introns characteristics in terms of length, GC content, maximum entropy at their extremity and conservation scores.

Table S7 | List of primers for qPCR validation of aberrant intron retention.

Table S8 | List of splicing factors used for Figure 1.

